# S-nitrosoglutathione Inhibits the Growth of Androgen-Receptor Dependent and Castration Resistant Prostate Cancer by Modulating FOXM1 Signaling

**DOI:** 10.1101/2022.08.21.504703

**Authors:** Anastasia Vedenko, Khushi Shah, Derek Van Booven, Fakiha Fridaus, Deepa Seetham, Himanshu Arora, Joshua M. Hare, Ranjith Ramasamy

## Abstract

Castration resistant prostate cancer (CRPC) is an advanced form of prostate cancer associated with loss of androgen receptor activity and resistance to AR-targeted therapies. CRPC is often associated with higher metastasis, invasion, and proliferation. Thus, there’s a strong need for additional molecular markers in CRPC, and therapies that target them directly. Transcriptomic profiling of prostate cancer patients revealed an upregulated signature specific to high-grade tumors in cell cycle progression, E2F targets, G2M checkpoint, DNA repair, Aurora Kinase B, and FOXM1 pathway. Validation in a separate cohort of neuroendocrine prostate cancer patients, the most resistant and invasive form of CRPC which lacks AR expression, showed CCP signature as a dominant driver gene set for NEPC. Moreover, high expression of mRNA targets in each of these pathways is associated with poor overall survival in NEPC patients. Gene set variation analysis revealed a strong inverse correlation between AR signaling and CCP.31, set of 31 markers associated with high proliferation, and a positive correlation between CCP.31 and NEPC signature. FOXM1, a prioritized candidate in CCP is shown to be upregulated in CRPC cell lines and human clinical sections of Gleason 9 patients. GSNO, a nitric oxide donor, known for its inhibitory effect on cell proliferation and an effective inhibitor of cell cycle checkpoints, was able to reverse CCP signature both in-vitro and in-vivo. GSNO exhibits a dependence on AR signaling in-vitro by showing mild effect on CRPC clones, but successfully inhibits tumor growth of AR-null CRPC xenografts. FOXM1 is also targeted by GSNO in-vivo and in-vitro models of both CRPC and AD prostate cancer. This establishes an important link between androgen receptor status in prostate cancer and proposes nitric oxide donors as therapeutic intervention. GSNO is effective against AR-null castration resistant prostate cancer.

**Significance:** Nitric oxide donor, S-glutathione-transferase inhibits tumor growth of castration resistant prostate cancer, regardless of androgen receptor expression status by targeting FOXM1, a critical regulator of cell cycle progression.

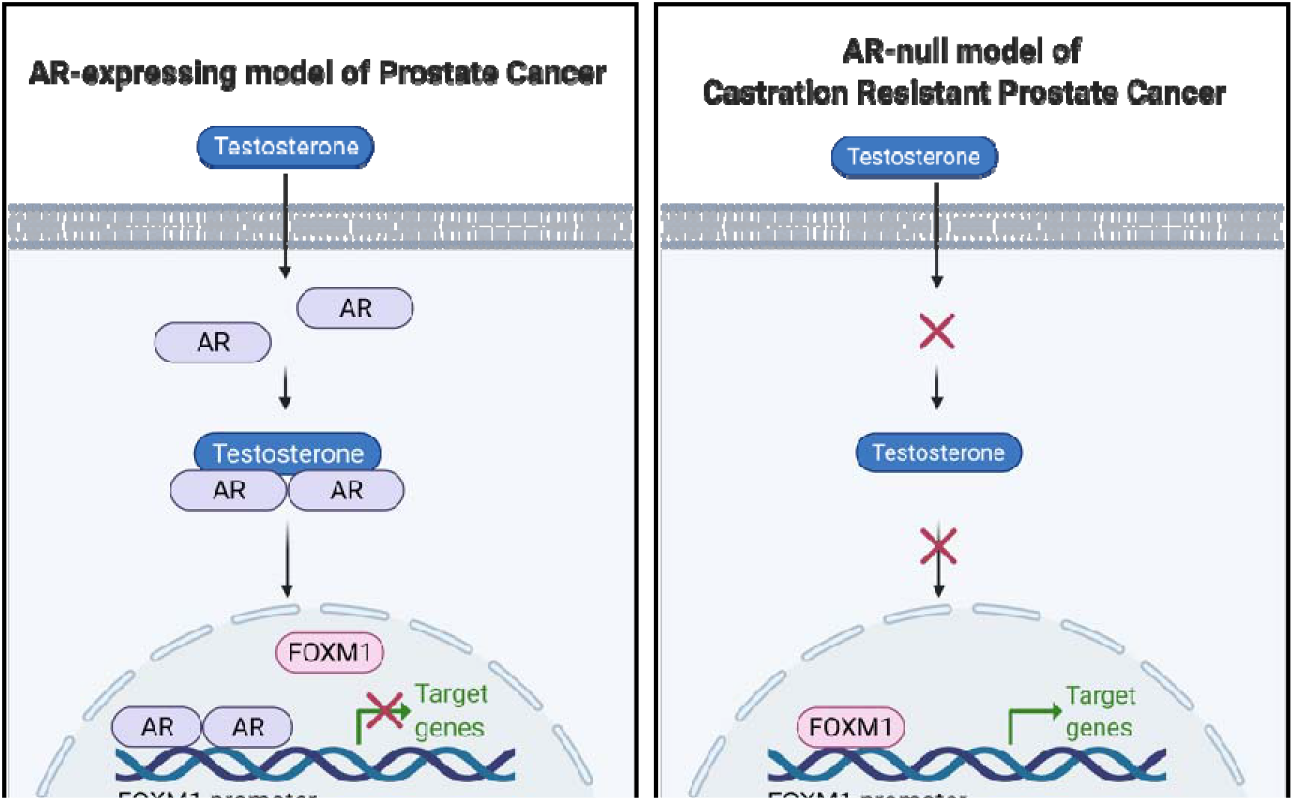

## Introduction

As early stages of prostate cancer rely on androgen receptor signaling (1), clinical intervention with androgen deprivation therapy (ADT) in order to reduce AR signaling, is effective at controlling tumor growth rate and improving patient outcome. However, while initially effective, these therapies ultimately fail, resulting in a rapid increase of prostate specific antigen (PSA) and emergence of a resistant state, known as castration resistant prostate cancer (CRPC). Therapies for CRPC are limited, and if left untreated, may progress to phenotypes with small cell-like or neuroendocrine features, which are highly invasive and can rapidly metastasize (11).

Accentuated by the fact that CRPC does not rely on AR, new molecular targets are being explored as well as therapeutics that can successfully target them. Section cores from malignant CRPC tumors have recently shown that patient samples often exhibit either a complete loss of AR expression or maintain a heterogenous level of AR, with some CRPC tumors continuing to express AR (2). While AR-expressing cells are still susceptible to second-line regimens by AR antagonists such as enzalutamide, AR-null cells show resistance de-novo, as well as a shorter cell cycle transit time, faster division, and preferred competitive advantage over AR-expressing cells in androgen depleted conditions (2).

Transcriptomic analysis of AR knockout mutants has revealed candidate programs which help maintain growth and promote survival of these clones, some of these include the FGF/MAPK pathway (3), stem cell associated genes (2), forkhead box M1 pathway (4), as well as multiple cell-cycle progression genes. A simple treatment of androgen-responsive clones with enzalutamide or a synthetic androgen inhibits or activates, respectively, E2F targets and G2M checkpoint transition, among others (5). Transcriptional profiling of CRPC patients showed an enrichment for the lethal and aggressive prostate cancer signature 1, PCS1, in patients with high Gleason score, which is mainly comprised of cell cycle genes (4).

The main objective of the present study was to identify and target signaling pathways in high-grade prostate cancer. We accomplished this surveying transcripional profiles of CRPC tumors from The Cancer Genome Atlas (TCGA) and Weill Cornell Medicine neuroendocrine prostate cancer patient cohort (2011-2015). Here we demonstrate the relevance of E2F targets, Aurora kinase B, G2M checkpoint, FOXM1 pathway, and cell cycle progression in high-grade tumors. Through survival analysis in NEPC dataset, we show that many gene targets within these pathways are overexpressed in NEPC patients and contribute to poor overall survival.

A group of therapeutic drugs known as nitric oxide donors have been widely explored as anti-cancer therapeutic agents. Their association with low toxicity, rapid metabolism, short half-life, and ability to spontaneously release free radical NO. moiety under physiological conditions has made NO donors attractive therapies in targeting tumor biology (6). Most commonly used NO donors are S-nitrosothiolos, nitro-vasodilators, and NONOates (6), of which S-nitrosothiols have shown to inhibit cell cycle and cell proliferation in solid tumors. GSNO has been particularly useful against CRPC (7), however its effect on AR-null clonotypes has not been widely explored. Utilizing different models of CRPC, in-vitro and in-vivo assays, we show that GSNO targets cell proliferation in an AR-dependent manner, while successfully inhibiting tumor growth of CRPC xenografts, regardless of their AR-status. This work has the potential to propose new therapies for the treatment of castration resistant prostate cancer.

## Materials and Methods

### Transcriptomic analysis of RNA-seq data

Tumor and nontumor (normal) samples were obtained from TCGA, downloaded through Firehose hosted by the Broad GDAC Data Portal, and Weill Cornell NEPC (2011-2015) (11–12). Trimmed reads were aligned to the hg19 human reference genome. Differential expression analysis was performed with edgeR package in Bioconductor. Clustering of tumor and nontumor samples was confirmed by principal component analysis. AR activity score was calculated using gene set variation analysis (GSVA) of curated signature of ten genes, ARG. 10, that are direct targets of AR (3). To classify pathways as significantly enriched with respect to AR activity, we applied two filtering criteria, including (i) ARG. 10 GSVA score, with “high-ARG. 10”>0 and “low-ARG.10”<0 and (ii) histological classification of patient biopsies with CRPC based on gleason score, such as gleason 8/9 patients were categorized as CRPC and gleason 6/7 as non-CRPC. For pathway identification, we used gene set enrichment analysis (GSEA) based on all expressed genes with an assigned annotation status. Pathways that reached significant false discovery rate (FDR) and normalized enrichment score (NES) of ± 2 were considered for further analysis. For correlation analysis, GSVA was used to calculate pathway score and Pearson method was used to perform correlation based on pathway-level rather than gene-specific. Kaplan-Meier curves were constructed using survival data from Cornell NEPC dataset for specific targets in enriched pathways, using “survival” package in R, and statistical significance was determined using nonparametric log rank test. Gene prioritization was done with a PubMed search on all candidates that passed Kaplan-Meier statistical significance threshold, set at 0.05, and only those candidates were included that had notable literature in tumor biology and for which antibodies are commercially available.

### Cell culture, cell lines, and reagents

LNCaP, 22Rv1, and H660 cells were purchased directly from American Type Culture Collection (ATCC). LNCaP-shAR and LNCaP-APIPC cells were donated by Dr. Peter Nelson of Fred Hutchinson Cancer Research Center and University of Washington, Seattle. Cells were maintained in RPMI-1640 supplemented with 1% Penicillin/Streptomycin solution (10,000 Units/ml Penicillin, 10,000 μg/ml Streptomycin) and 5% fetal bovine serum (FBS) for LNCaP and 22Rv1 or 5% charcoal stripped FBS for H660 cells. Reagents studied here and included in the experiments were S-nitrosoglutathione (Cayman 82240), JS-K (Chem Cruz sc-211683), Enzalutamide (Selleckchem S1250), and 5α-Dihydrotestosterone (DHT) solution (Sigma-Aldrich D-073-1ML).

### Cell fractionation for RT-qPCR

Adherent and non-adherent cells were dissolved with TrypLE Express solution (Gibco 12604-021) after washing twice with DPBS. Cells were centrifuged at 4°C and 1250rpm for 5 minutes, after which the pellets were washed twice with chilled DPBS. Pelleted cells were finally resuspended in TRI Reagent (Sigma T9424-200ML) with multiple pipetting, followed by RNA extraction according to manufacturer’s protocol instructions. RNA quality was assessed by 260/280 ratio on spectrophotometer, and reverse transcribed to cDNA using High Capacity cDNA Reverse Transcription Kit (Applied Biosystems 4368814) following manufacturer’s recommendations. RT-qPCR was performed using SsoAdvanced SYBR Green Supermix (Bio-Rad 1725271). A complete list of sequences used in this study are reported in Summary Table S1.

### Cell fractionation for Western blotting

Following similar extraction protocol for RT-qPCR, final cell pellets were instead resuspended in RIPA buffer supplemented with protease inhibitor cocktail (Sigma SRE0055-1BO), and following a brief incubation period on ice, the cells were sonicated for at least 30 seconds in three consecutive bursts. The lysed cells were then centrifuged at 10,000 and 4°C for 10min, and clear supernatant was collected for immunoblotting analysis. Immunoblotting was performed using antibodies against AR (Santa Cruz Biotechnology sc-7305), PSA (Santa Cruz Biotechnology sc-7316), FOXM1 (Santa Cruz Biotechnology sc-376471), CDC20 (Santa Cruz Biotechnology sc-13162), TMPRSS2 (Santa Cruz Biotechnology sc-515727), FKBP51 (Santa Cruz Biotechnology sc-271547), BUBR1 (Santa Cruz Biotechnology sc-47744), NKX3.1 (Santa Cruz Biotechnology sc-393190), TOPOIIa (Santa Cruz Biotechnology sc-365916), CENP-E (Santa Cruz Biotechnology sc-376685), and GAPDH (Invitrogen ZG003).

### MTT assay

Cell proliferation was measured in a 96-well format. Cells were seeded at 5,000 cells per well, after 24 hours, spent media was collected and treatment was initiated. Measurements were taken every 24-48 hours by incubating the cells with MTT reagent A (Millipore CT01-5) for 2-4 hours followed by DMSO for 10 minutes. Calorimetry measurement was recorded at 562nm on a micro-plate reader SpectraMax M5 (Molecular Devices).

### Migration assay

Cells were seeded at confluence 8×10^5^ (LNCaP) or 1×10^6^ (shAR, APIPC) in a 6-well plate. After 24 hours, the cells were gently washed and scraped directly diagonally across with 20ul tip to leave an indentation and allow a clean space for cell migration. Cells were imaged every 24-48 hours, and migration distance was measured in ImageJ by subtracting the reference (zero time point of the corresponding cell line). Statistical analysis was performed in GraphPad Prism software.

### Apoptosis assay

Cells were seeded in duplicates and treatment was initiated after 24 hours. Staining was performed with Annexin V kit following manufacturer’s recommendations (Thermo Fisher Scientific). Necrotic and apoptotic cell populations were counted every 24 hours for 3 consecutive days by FACS sorting.

### IHC

Tissue slides were deparaffinized, processed for antigen retrieval, and blocked with appropriate reagents in the HRP/DAP detection IHC kit (abcam ab64264) following manufacturer’s protocol, after which primary antibodies were applied for FOXM1 (1:50, Santa Cruz Biotechnology sc-376471), AR (1:50 Santa Cruz Biotechnology sc-7305), PSA (1:50 Santa Cruz Biotechnology sc-7316), and Ki67 (1:50 Invitrogen). Following secondary antibody incubation, tissue slides were imaged, and staining intensity of at least 16 different sections was averaged and quantified with ImageJ. Statistical analysis was performed in Prism software.

### Castration, xenografting, treatment, and tumor size measurement

NOD-Cg Prkidc^scid^ mice were purchased from The Jackson Laboratory. Castration was performed on two groups of mice receiving APIPC or shAR xenografts by a qualified surgical technician, following guidelines outlined in protocol no. 17-113, approved by Institutional Animal Care and Use Committee (IACUC) at the University of Miami Miller School of Medicine, FL. After 7-days post-castration, xenografts were injected subcutaneously in the thoracic cavity of the animal, using 5×10^6^ LNCaP, shAR, or APIPC cell suspension, diluted 50:50 in Matrigel. The mice were then further divided into two groups, and treatment with GSNO or PBS was initiated using 10mg/kg/day by injection in the Intraperitoneal (IP) cavity, with daily injections allowed to proceed for two weeks. Tumor growth was monitored twice a week for 60 days, and tumor volume was calculated using the formula V=(LxW^2^)/2.

### Statistical analysis

Statistical analysis was performed in GraphPad Prism 9. Two-way ANOVA or a paired t-test were used as appropriate. A p-value of 0.05 was considered to be significant. Data are reported as mean ± SEM of technical triplicates.

## Results

### Transcriptomic analysis of castration resistant prostate cancer is characterized by a marked reduction in AR activity in high-grade tumors

Using publicly available RNA-seq data from the cancer genome atlas (TCGA), we first sought to establish if there was a correlation between androgen receptor expression and grade of cancer in prostate adenocarcinoma and their un-matched normal controls. mRNA expression of AR gene alone, quantified through RNA-seq, did not reach significance in adenocarcinoma tumors when compared to normal samples. This is not surprising, given that AR might not be active, yet maintain its expression status through truncated splice variants in CRPC. Therefore, we took advantage of a gene signature AR.10, which interrogates many of the downstream targets of AR and can be used to interrogate AR activity as a whole in a particular sample. Adenocarcinoma samples showed an uncharacteristic upregulation of many AR.10 targets (ALDH1A3, KLK2, KLK3, NKX3.1, PMEPA1, STEAP4) in tumor samples (**Supplementary Fig. S1A-B**). This could be explained by two scenarios. The normal samples are mainly representative of biopsy patients with Gleason 7 tumors of the surrounding non-neoplasm. Second, adenocarcinoma group represents a heterogeneous population of patient biopsies ranging from primary prostate low or medium-grade cancer (Gleason 6 and 7) to castration resistant high-grade prostate tumors, often associated with Gleason 8 and above. Thus, further combining patient groups into Gleason 6/7 and Gleason 8/9 showed a significant downregulation of AR.10 in high-grade tumors (KLK2, KLK3, PART1) (**Fig. 1E**). This suggests a downregulation of AR activity in advanced prostate cancer. Hierarchical analysis revealed two clusters, with cluster 2 mainly comprised of Gleason6/7 patients and showing high expression for androgen response genes in unsupervised (**Fig. 1B**) and supervised (**Fig. 1C**) hierarchical clustering. To understand if AR activity correlated with Gleason score, we used gene set variation analysis (GSVA), curated with AR.10 gene set. Increase in Gleason score showed a very strong and positive downregulation (Kruskal-Wallis, p=5.4e-14) of AR.10 score between Gleason 6 and Gleason 9 tumors (**Fig. 1D**). This further validates that AR activity is important for progression to CRPC. Gene set enrichment analysis (GSEA) of Gleason 8/9 tumors showed an enrichment for Hallmark Androgen Response (NES=1.45, FDR=0.036), Reactome stimulation of androgen receptor regulated genes KLK2 and KLK3 by PKN1 (NES=-2.756, FDR=0.0) in adenocarcinoma, TCGA, Nelson response to Androgen_UP (NES=-2.638, FDR=0.0), (**Supplemantary Fig. S1C-D**) and DOANE response to Androgen_UP (NES=-2.40, FDR=0.0) in neuroendocrine prostate cancer (Weill Medicine Cornell RNAseq dataset). All together, these results show a reduction in AR activity in CRPC, thus allowing us to interrogate this feature for downstream analysis of pathways specific to clinically relevant signatures of the most lethal and aggressive form of AR-negative CRPC.

**Figure 1.**
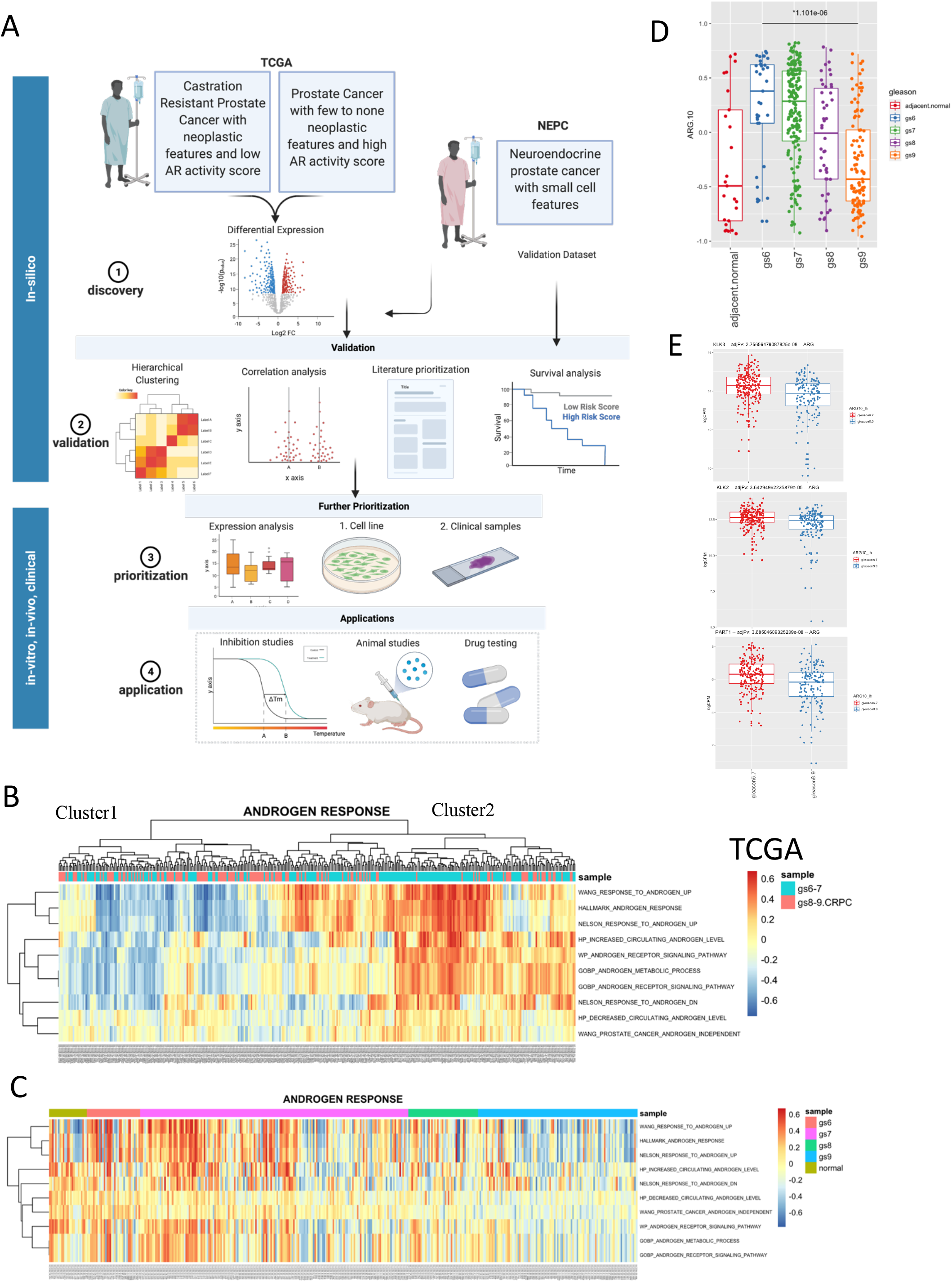
CRPC is associated with a reduction in AR activity in high-grade tumors. **A,** pipeline of the steps taken to select candidates highly expressed in CRPC. **B,** Hierarchical clustering analysis by androgen-specific MSigDB gene sets reveal two clusters. Cluster 1 is enriched for CRPC, gleason8-9 patients, while cluster2 is enriched for low-grade tumors, gleason6-7. **C,** Supervised un-clustered heatmap of androgen response gene sets, ordered by increasing gleason score. Far right corresponds to gleason-9, far left to normal samples. **D,** Gene set variation analysis for AR.10 gene signature (p=1.1e-6, student’s two-tailed t-test). **E,** Expression of significantly downregulated AR.10 targets in gleason score 8-9 patients. Adjusted p-value is displayed above the plots, adjusted by the number of candidates within the pathway and patient samples in TCGA (n=495), significance determined by two-tailed, student’s t-test. MSigDB-GSEA gene sets chosen at random as a sampling for androgen receptor association and does not include all possible signaling targets of androgen response. TCGA, the cancer genome atlas; ARG, androgen receptor genes; gs, gleason score.

### Transcriptomic analysis of high-grade, low AR-activity prostate tumors is enriched for cell cycle progression genes

We wanted to investigate what programs drive and sustain CRPC when it is devoid of androgen receptor expression. For this, we further interrogated RNA-seq data from the TCGA to gain insight into transcriptional profile of CRPC patients (**Fig. 1A**). Utilizing GSVA scoring system for AR. 10 signature, we divided the cohort into two modules, low or high AR. 10 score-expressing patients. To make the analysis even more clinically-relevant and filter out potential confounders (ie. patients with uncharacteristically low-expressing AR.10 that do not have high grade cancer), we further selected only those patients in high Gleason (Gleason8/9) with low AR.10 and defined them as “high-grade AR-low CRPC”. Gene set enrichment analysis revealed that patients in AR.10-low group are enriched for MSigDB gene sets involved in cell proliferation: cell cycle checkpoint, aurora B pathway, ATR pathway, E2F targets, G2M checkpoint, PLK, FOXM1 pathway, and oncogenic signatures, MYC targets and P53 regulation in TCGA (**Fig 2A**) in NEPC (**Fig. 2B**). A previously published signature (5) of cell cycle progression CCP.31 was also included in hierarchical clustering analysis, and showed a similar pattern of clustering to the cell cycle pathways identified here, and was thus included in downstream analysis of gene candidates. Supervised hierarchical clustering of these pathways across all TCGA samples revealed two main clusters: cluster 1 was mainly dominated by ARG.10-high-Gs6-7 patient samples, while cluster 2 was highly dominated by ARG.10-low-Gs8-9 patients samples (**Fig. 2C**). Cluster 2 was associated with statistically significant upregulation of CCP.31 genes (p=6e-4). A validation dataset was used from Weill Medicine Cornell neuroendocrine prostate cancer patients similarly stratified into low/high groups by AR.10 score, showed a similar downregulation of CCP.31 genes in the low-AR.10 NEPC/CRPC patient category (p=3e-6) (**Fig. 2D-E**). Unsupervised hierarchical clustering of 31 genes in CCP similarly showed two distinct clusters, where high expression of all 31 genes was mainly observed in high-risk CRPC patients (**Supplementary Fig. 2SA**). Therefore, these genes were prioritized for further analysis. We found that ARG.10 signature has an inverse correlation with CCP.31 signature by regression analysis (R=-0.38, p=8.9e-11), and that CCP.31 signature has a positive correlation with neuroendocrine prostate cancer gene signature NEPC.10 (3), (R=0.2, p=0.00085) (**Supplementary Fig. 2SC**). This suggests the possible regulatory control of CCP genes by androgen receptor signaling, as well as establishes their importance in high grade AR negative prostate cancers. Principal component analysis showed a separate cloud for normal samples apart from adenocarcinoma samples (**Supplementary Fig. 2B**). High expression of these markers was associated with Adenocarcinoma (**Supplementary Fig. 2SD**) and Gleason8/9 (**Supplementary Fig. 2F**), and NEPC (**Supplementary Fig. 2E**). Furthermore, high expression of these markers was associated with poor overall survival in our NEPC validation dataset (**Fig. 2F**). We further prioritized these markers based on literature and antibody database search to select only those candidates that were associated with prostate or other cancers (4,8–10).

**Figure 2.**
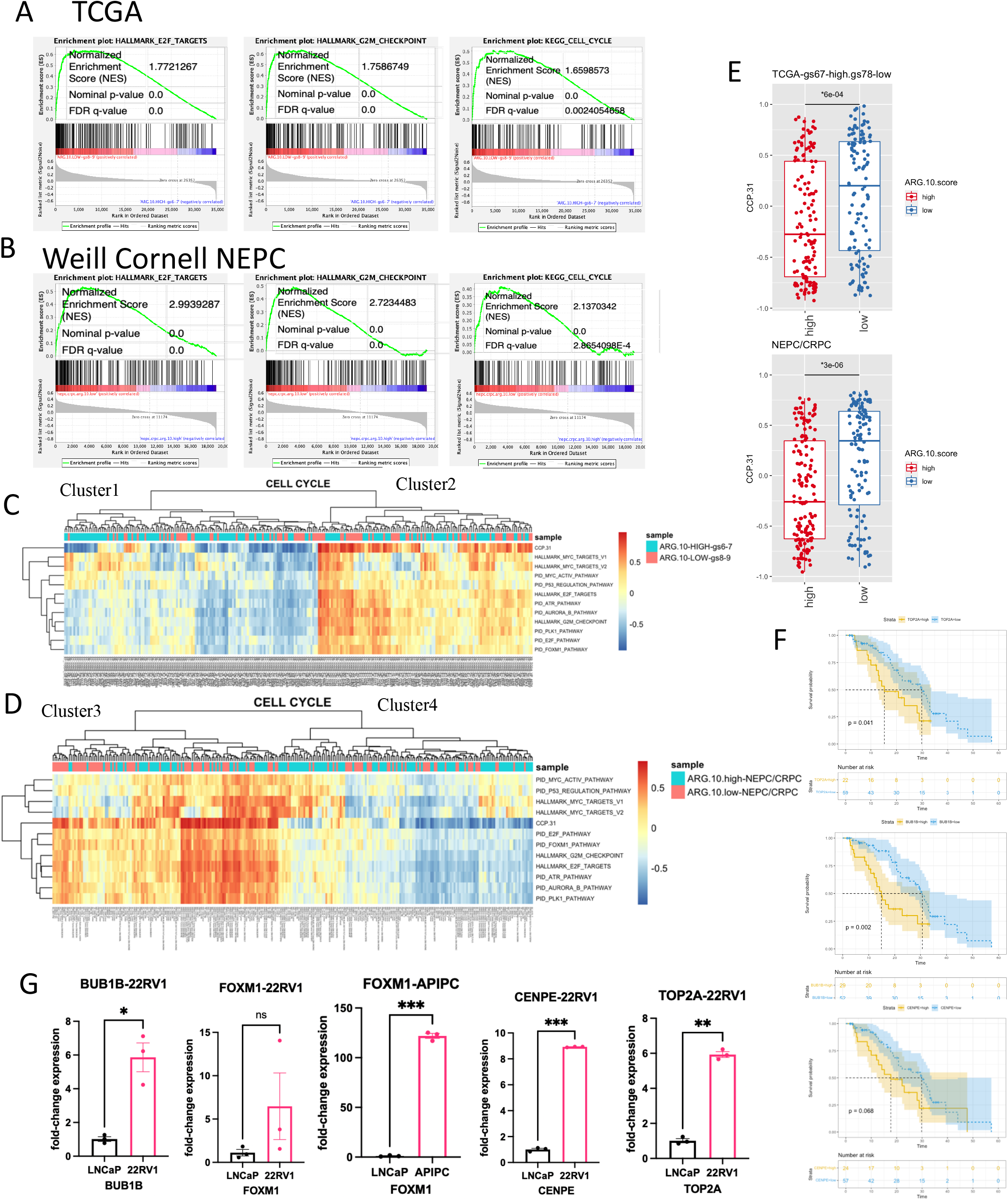
Transcriptomic analysis of high-grade, low AR-activity prostate tumors is enriched for cell cycle progression genes. **A-E,** Tumor samples were grouped into high (AR.10>0; n=214) and low (AR.10<0; n=176) AR expressing samples. Representative enrichment plots for TCGA, **A,** and Weill Medicine Cornell NEPC cohort, **B**. Hierarchical clustering analysis with two defined clusters, corresponding to sample categories indicated for MSigDB data sets from the top ten candidates in GSEA analysis for TCGA, **C,** and NEPC, **D**. **E,** boxplots of mRNA expression scored via gene set variation analysis using CCP.31 curated geneset in TCGA (top) and NEPC (bottom). *P-value denoted is calculated by analysis of variance (ANOVA). Thick middle bar of boxplot represents the median, top and bottom bars represent first and third quartiles. **F,** Kaplan-Meier overall survival of representative targets in CCP.31 gene list calculated from survival data of NEPC/Cornell patient dataset. Statistical significance was determined by log rank test and hazard ratio by Cox proportional hazards model. **G,** fold change expression of prioritized candidates from CCP.31 gene list in indicated CRPC cell line models. Statistical significance is determined by student’s t-test; ***, P<0.001; **, P<0.01; * P<0.05.

### Cell cycle genes are enriched in advanced prostate cancer cell lines and Gleason score-9 clinical samples

We further validated shortlisted candidates in castration resistant prostate cancer, 22Rv1 and neuroendocrine prostate cancer cell line, H660. Both of these in-vitro models showed a higher than normal expression of CCP.31 genes, particularly FOXM1, BUB1B, CENPE, and TOP2A based on prioritization criteria. To validate the hypothesis that androgen receptor loss is associated with higher expression of these genes, we utilized Androgen-Receptor program independent prostate cancer cell line (3), APIPC. mRNA expression of FOXM1 was significantly upregulated in APIPC and 22Rv1 cells (**Fig. 3A**). We further treated androgen dependent prostate cancer cells, LNCaP, with Enzalutamide, a direct AR-antagonist, to further test our hypothesis and found that expression of CENPE was significantly upregulated in Enza treated cells, while FOXM1, BUB1B, and TOP2A showed a similar pattern of inhibition (**Fig. 3B**). We next evaluated the expression of FOXM1 in Gleason 9 patient sections and non-matched normal prostate controls and found that FOXM1 expression was significantly upregulated in tumor sections. These results suggest that cell cycle progression is activated in CRPC, and thus we focused on FOXM1 for further analysis.

**Figure 3.**
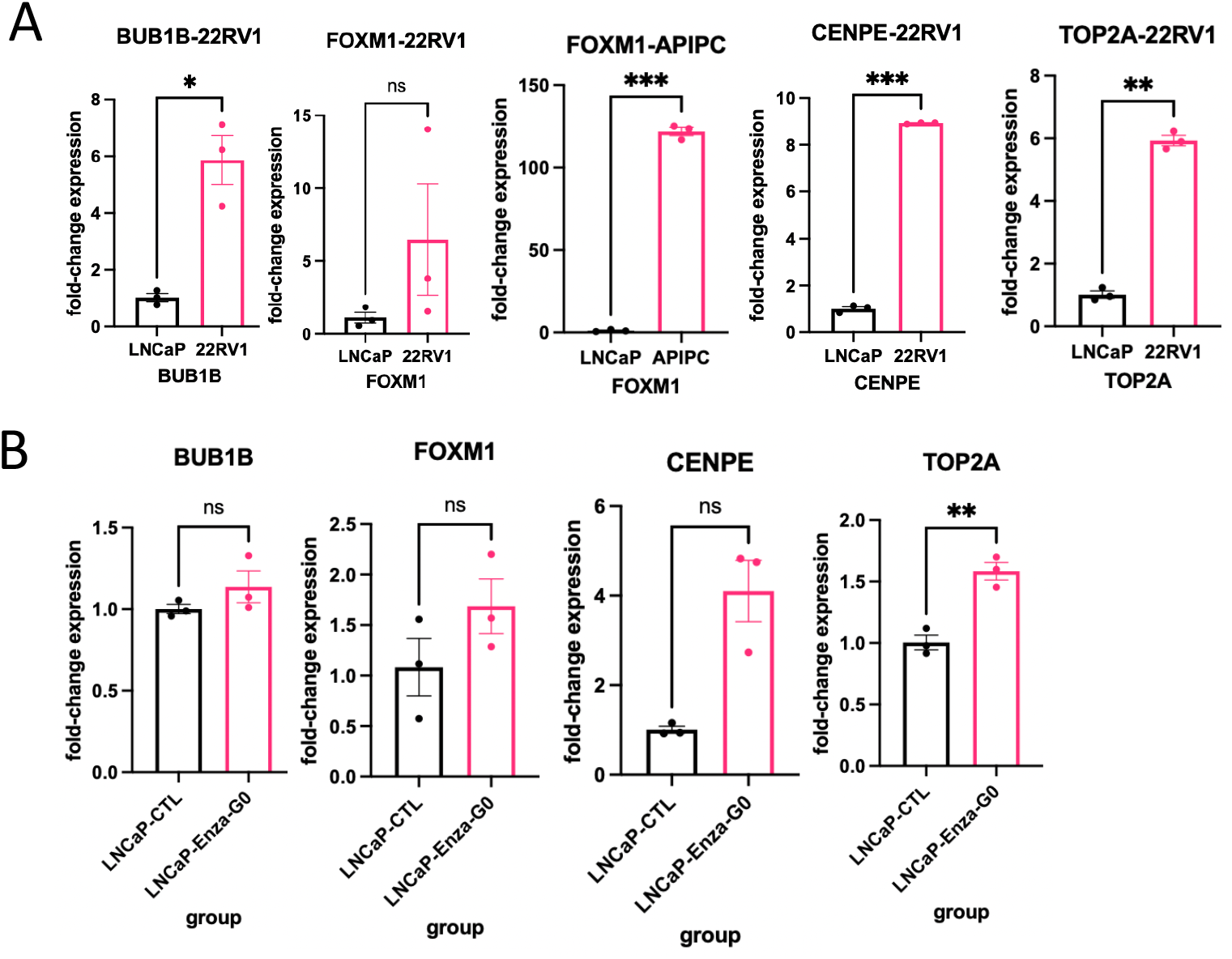
Cell cycle genes are enriched in advanced prostate cancer cell lines and gleason score-9 clinical samples. **A,** q-PCR analysis of fold change expression of corresponding cell line relative to expression in LNCaP, which as defined as 1.0. Fold-change expression of each pair of samples was derived by the formula 2^-ΔCt^ [ΔC_t_=C_t_(gene)-ΔC_t_(GAPDH)]. X-axis indicates fold-change of each gene expression in corresponding cell line relative to LNCaP (defined as 1.0). **B,** qPCR analysis of expression of representative genes in LNCaP treated with 4μM Enzalutamide. Fold-change expression is reported relative to LNCaP vehicle (DMSO) treated samples. Data are reported as mean ± SEM of technical triplicates from a representative experiment (student’s t-test). ***, P<0.001; **, P<0.01; *, P<0.05; ns, not significant.

### Inhibition of cell proliferation by nitric oxide donor, GSNO, depends on androgen receptor expression status in prostate cancer

We next tested the ability of a nitric oxide donor, GSNO, on proliferation capacity of prostate cancer cells with variable degrees of androgen receptor expression status. We (7) and others have shown that GSNO is able to inhibit the proliferation of 22Rv1 cells in vivo and in vitro, however 22Rv1 model which was used in that study has a transient expression of AR. Here, we wanted to test the hypothesis that GSNO is AR independent and can inhibit the growth of AR-negative prostate cancer. APIPC cells and their parent derivative, LNCaP, were used for the study design. While cell proliferation was significantly reduced in LNCaP cells upon GSNO treatment (IC_50_=93μM), APIPC proliferation was unaffected during two-week inhibitor response (IC_50_=244μM), and intermediate AR-level LNCaP-shAR cells showed a similar curve (IC_50_=203μM) (**Fig. 4A-C**). Similarly, migration of APIPC cells was slower than LNCaP (**Fig. 4D-F**), and cell cycle by Annexin V stain of 22Rv1 and H660 cells showed a decrease in necrotic population only in LNCaP (74%) and not in 22Rv1 (100%) or H660 (125%) (**Fig. 4E**). This data suggests that GSNO targets prostate cancer in androgen receptor dependent manner, and patient tumors that exhibit complete loss of AR might not be susceptible to inhibitory effect by GSNO. We further recapitulated this model using AR agonist, 5α-Dihydrotestosterone (DHT) and antagonist, Enzalutamide. Upon GSNO in combination with Enzalutamide treatment, we observed no inhibition of cell proliferation of LNCaP cells by MTT assay (IC_50_=wide), yet a strong decrease in cell proliferation in combination with DHT (IC_50_=387μM) (**Fig. 5D-E**). Subsequent studies confirmed our observations, such as treatment with GSNO plus DHT on cell lines responsive to androgen stimulation, LNCaP-shAR, reduced IC_50_ value back to the original level observed in LNCaP (IC_50_=102μM) (**Fig. 5F-G**). This also suggests that GSNO is acting through an AR-specific mechanism.

**Figure 4.**
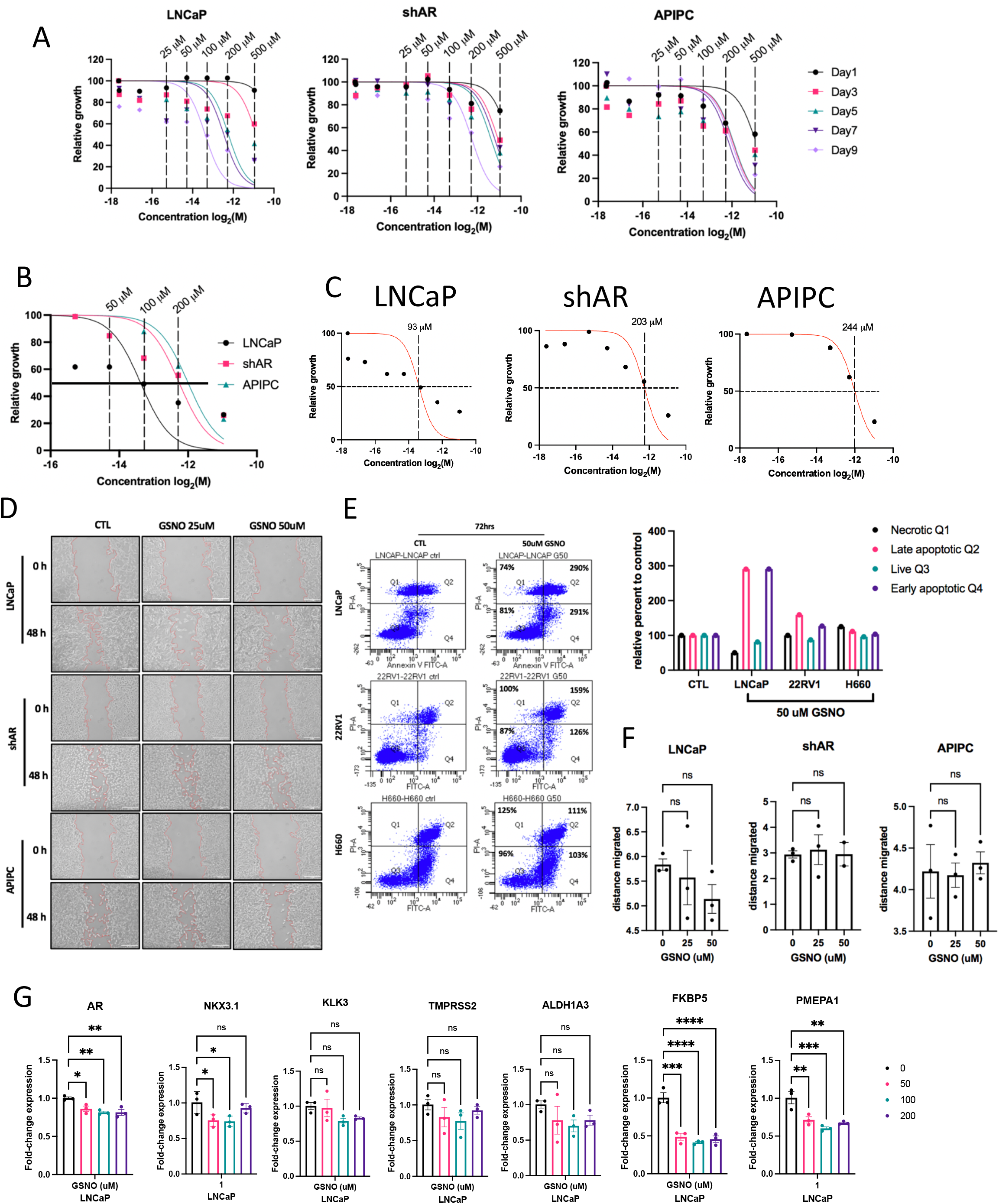
Inhibition of cell proliferation by GSNO depends on androgen receptor expression status in prostate cancer. **A,** cell proliferation curves of indicated cell line treated with GSNO at log2 - transformed concentration for indicated number of days. Relative growth (x-axis) is calculated in reference to vehicle-treated control and represents percent growth (out of 100%). Vertical dashed lines represent GSNO concentrations in micromolar. **B-C,** Proliferation curves of three indicated cell lines treated with increasing concentrations of GSNO. Dashed lines represent original GSNO concentrations for clarity, solid bar marks the location of the IC_50_, depicted for each CRPC cell line separately in **C**. **D,** migration assay of each indicated cell line treated with two concentrations of GSNO for 48 hours. Images were taken every 24 hours. Scale bar, 1μm. Images were taken with Nikon microscope. **E,** Annexin V FITC-A scatter of indicated CRPC cells treated or untreated with 50μM GSNO for 72 hours, with quartiles corresponding to necrotic, q1, late apoptotic, q2, live, q3, and early apoptotic, q4. Percentages of cells in each phase are depicted in quartile boxes above the scatter, calculated relative to each cell line’s corresponding untreated control. **F,** quantification and statistical analysis of migration assay in **D**. **G,** qPCR analysis of GSNO treatment on AR.10 genes. Data are reported as mean ± SEM of technical replicates (student’s t-test). ***, P<0.001; **, P<0.01; *, P<0.05; ns, not significant.

**Figure 5.**
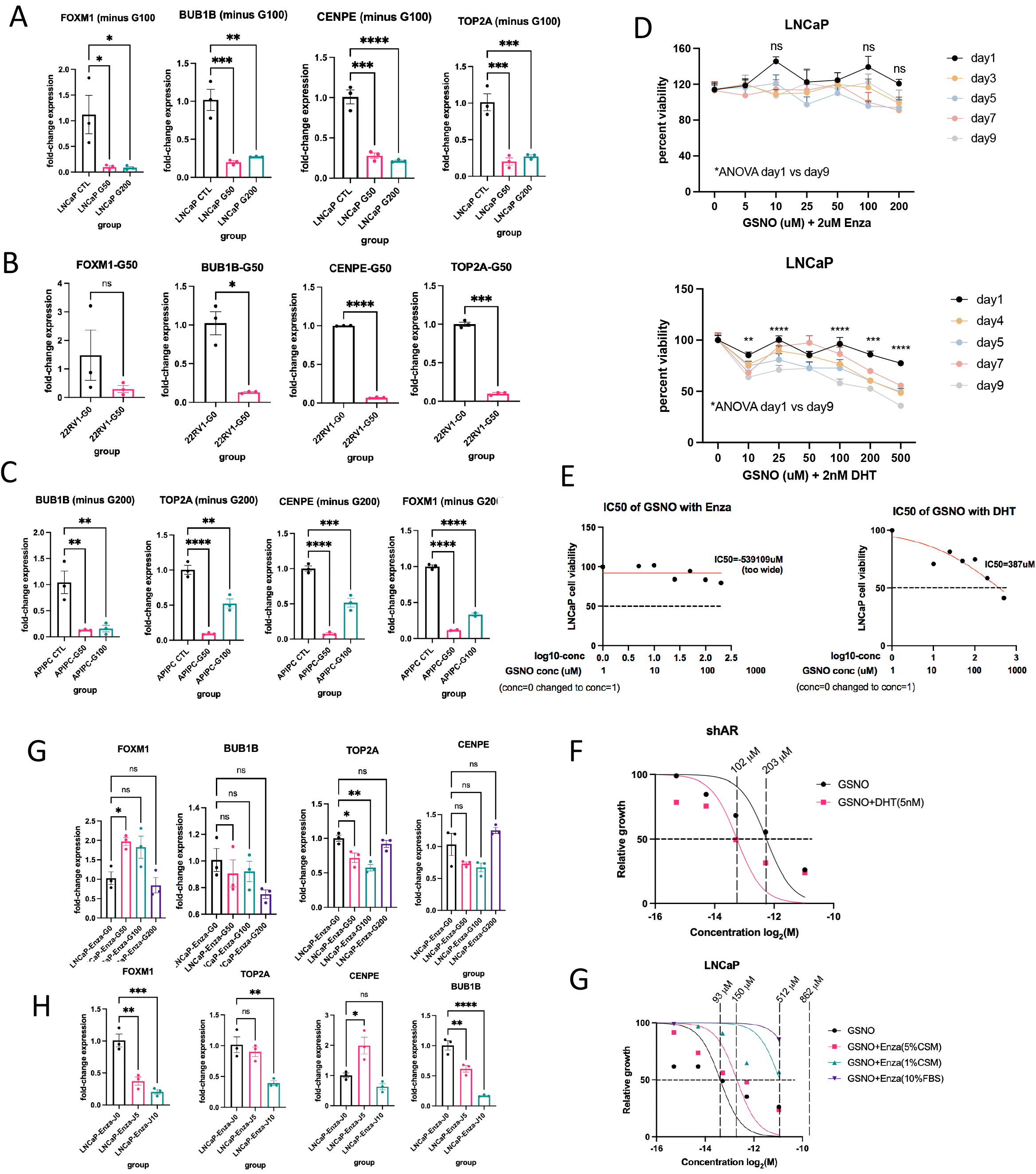
GSNO targets markers associated with CRPC. **A-C, G-H,** qPCR analysis of expression of representative genes from CCP.31 signature that reached significance in LNCaP, **A**; 22Rv1, **B**; APIPC, **C**; when treated with GSNO or untreated control. Expression was also assessed in LNCaP treated with 4μM Enzalutamide in combination with GSNO, **G**; or JS-K, **H**. Data are reported as mean ± SEM of technical triplicates (student’s t-test, ****P<0.0001; ***, P<0.001; **, P<0.01; *, P<0.05; ns, not significant). Fold change is calculated relative to untreated control in each cell line (defined as 1.0). **D,** Cell proliferation curves calculated out of 100% relative to untreated sample at each day the measurement was taken of LNCaP in combination with 2μM Enzalutamide (top) and LNCaP treated with GSNO in combination with 2nM DHT (bottom). **E,** IC_50_ values of two treatment conditions in **D** for GSNO+DHT (right) and GSNO+Enza (left). Dashed line defines location of the IC_50_ value for each experiment. **F,** proliferation curve of additional CRPC cell line, shAR, responsive to androgens. **G,** inhibitor plots with further extension of IC_50_ value in different FBS or charcoal-stripped FBS media, show a link between FBS concentration and androgen dependence on IC_50_. DTH, dihydrotestosterone; Enza, enzalutamide, GSNO, S-nitrosoglutathione.

### GSNO targets markers associated with CRPC

We next tested the ability of GSNO to inhibit markers associated with CRPC that were prioritized from transcriptomic analysis in our study. Treatment with GSNO at 50-100μM showed a statistically significant inhibition of FOXM1, BUB1B, CENPE, and TOP2A in LNCaP, 22Rv1, and APIPC cells by RT-qPCR (**Fig. 5A**) in LNCaP and to a higher degree in 22Rv1 (**Fig. 5B**) and APIPC (**Fig. 5C**). We also showed a similar pattern of inhibition by western blotting (FOXM1, CENP-E, BUB1B, CDC20 and TOPOIIa) in LNCaP. Furthermore, inhibition of AR with Enzalutamide in LNCaP cells significantly inhibited BUB1B, TOP2A, and CENPE (**Fig. 5G**). Treatment with a different group of nitric oxide donor, a diazeneiumdiolate prodrug, JS-K, produced similar decrease in FOXM1, TOP2A, and BUB1B (**Fig. 5H**). Furthermore, GSNO reduced majority of AR.10 pathway genes (AR, NKX3.1, FKBP51, PMEPA1, KLK3, TMPRSS2, and ALDH1A3) in LNCaP cells (**Fig. 4G**). These results suggests that GSNO inhibits cell cycle markers in cell line models of both AD PrCa and CRPC.

### GSNO reduces tumor burden in AR-null xenograft model of CRPC

GSNO was previously shown to reduce tumor burden of CRPC xenografts by our group (7). Therefore, in this study design, we tested if androgen receptor was the determining factor and if GSNO is able to target tumors that exhibit a complete loss of AR (**Fig. 6A**). Mice with APIPC xenografts responded similarly to GSNO treatment in both LNCaP and shAR xenografts with a notable reduction in tumor volume, observable just two weeks post-cessation of treatment. By the end of 8-week period, tumor volumes were almost undetectable in GSNO-treated shAR animals and remained low, near 200 cubic centimeters in LNCaP and APIPC (**Fig. 6B-C**). Reduction in expression of AR.10 markers (AR, TMPRSS2, NKX3.1, FKBP5, and ALDH1A3) was detected by RT-qPCR in LNCaP GSNO treated mice (**Fig. 6D**). GSNO also reduced CCP pathway genes (FOXM1, CENPE, TOP2A) in APIPC (**Fig. 6F**) and LNCaP (FOXM1, CENPE) (**Fig. 6E**) by RT-qPCR. This suggests a separate role of GSNO and variability of its targeting based on the microenvironment context.

**Figure 6.**
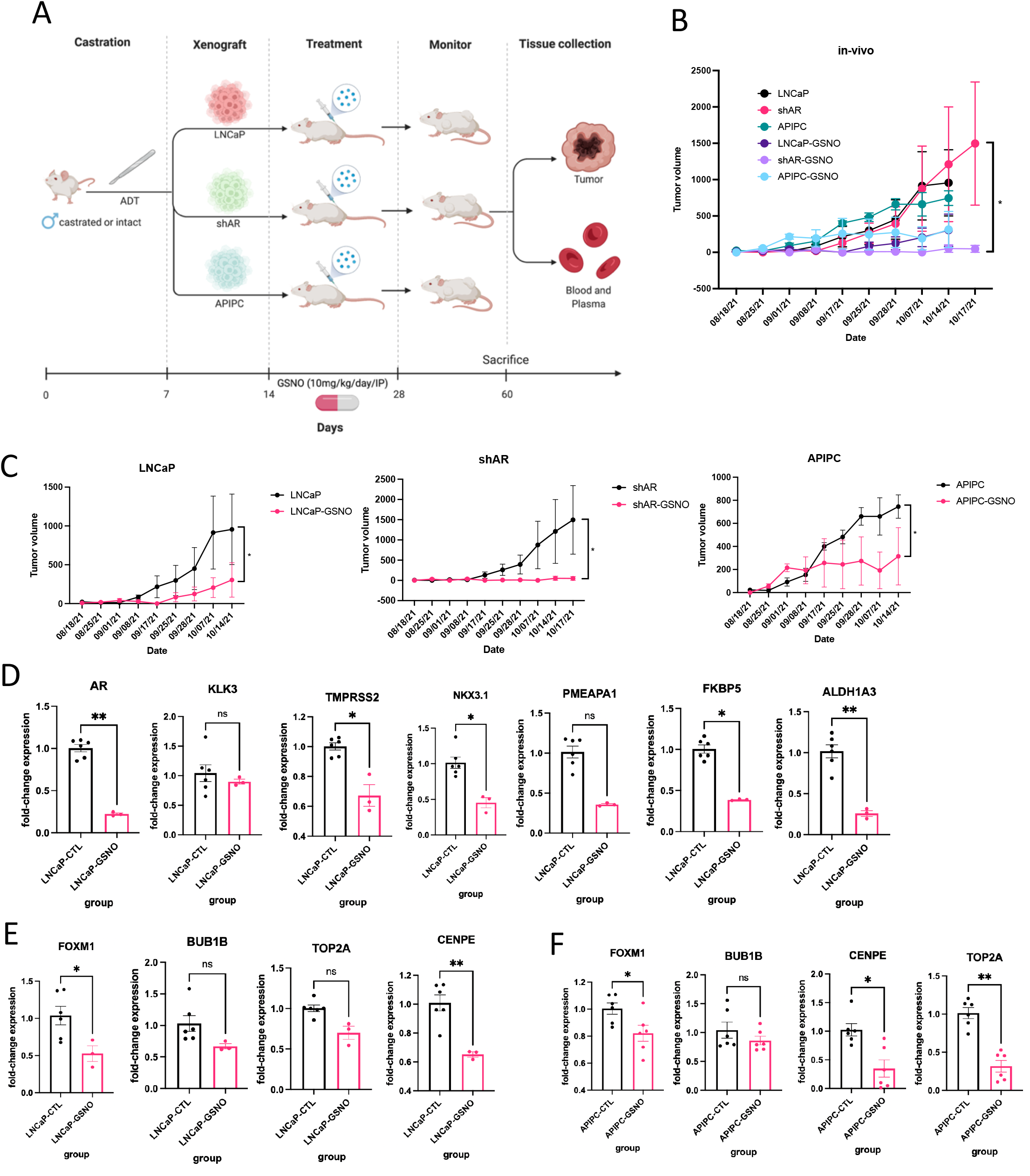
GSNO reduces tumor burden in AR-null xenograft model of CRPC. **A,** study design and timeline of procedures and animal groups used for this study. Tumor volume in units cc (cubic centimeter) calculated using the formula V=(LxW^2^)/2 for all assessed models, **B,** and each cell line separately, **C**. Signficance is determined by ANOVA, **B,** or Student t-test, **C**. **D,** qPCR analysis of ARG.10 signature genes reduced by GSNO treatment in xenografts of LNCaP animals, relative to the untreated control group, LNCaP, expressed as fold-change, set at 1.0. **E-F,** q-PCR of expression of prioritized markers in CCP.31 gene set that were significantly reduced by GSNO treatment in xenografts of animals with LNCaP, **E**, or APIPC tumors, **F**. Data are reported as mean ± SEM of technical replicates (student’s t-test, ***, P<0.001; **, P<0.01; *, P<0.05; ns, not significant). ADT, androgen deprivation therapy.

## Discussion

Major cause of cancer related death from castration resistant prostate cancer continues to dominate in western men. Therapies for CRPC are extremely limited, and while tumor clone reliance on AR signaling is targetable by AR antagonism, clones with complete absence of AR are often associated with greater plasticity, self-renewing potential, and invasive properties (1-3). Thus, development of additional therapies for AR-negative prostate cancer subtypes are extremely and urgently needed. In this study, we transcriptionally profiled the cancer genome atlas prostate adenocarcinoma RNA-seq data and a validation dataset of neuroendocrine prostate cancer patients, to design one of a kind study, where androgen receptor activity was used as a predictor of high-risk CRPC patients. By doing it this way, we were able to tease apart only those patients with lowest level of AR activity while maintaining a high-grade tumor. Other studies have similarly profiled RNA-seq datasets, however using more conventional methods of group assignment such as Gleason score or cancer grade to predict molecular pathways (4). This study design allowed us to reach normalized enrichment score, NES of 3.25 for androgen response by GSEA, thus improving confidence in our methods. Applying GSVA scoring in our analysis for AR signaling pathway, we found that AR activity was clinically relevant, as Gleason 6 patients exhibited AR activity of 0.5 (GSVA scale: −1 to +1) and Gleason 9 GSVA score was close to −0.5. Furthermore, adenocarcinoma patients had a much more increased GSVA score (positive) than normal patients (−0.5). We found many significant pathways enriched by AR activity score in patients with CRPC, several of which have been previously published (4), but we also identified new relevant pathways that we believe could shed light on mechanisms behind resistance development when AR status alone cannot be used as a predictor of progression (E2F targets, ATR, Aurora B kinase, G2M targets, and PLK1 pathway). Our validation dataset on neuroendocrine prostate cancer patients confirmed many of these findings. Moreover, specific gene targets were identified by survival analysis, which showed that their higher expression was associated with poor overall survival. These markers should further be evaluated for potential targeted and precision therapies for patients with resistant tumors and low AR-status. We were also the first to show that cell cycle progression signature has an inverse correlation with AR activity score and a positive correlation with neuroendocrine differentiation. This strengthens our belief that targeted therapies against CRPC should be aimed at cell proliferation and cell cycle checkpoints. We also validated that FOXM1, a top marker on our list, by literature and database prioritization, is indeed highly expressed in CRPC and NEPC patient cell lines by RT-qPCR as well as patient Gleason 9 clinical samples by IHC, in accordance with previously published reports (4, 8–10). This further increases our confidence that our methods are robust. We decided to focus on nitric oxide donor, GSNO, as a therapeutic intervention in AR-low cancers because it was previously shown to be efficacious in CRPC patient cell line and xenograft models by significantly reducing cell proliferation and tumor burden (7) as well as its unique ability to be a valuable therapeutic in cell cycle and proliferation studies. We were able to show that while the drug is ineffective against AR-low cancers in-vitro by MTT, migration, and was associated with low percentage of necrosis in Annexin V assay, it is extremely effective against AR-null cancers in-vivo by reducing tumor burden significantly only within the first two weeks post-treatment. Other groups have shown that GSNO is effective against LNCaP, 22Rv1, and several forms of small hairpin targeted AR models, but this study is the first of this kind to show that GSNO is effective against AR-null prostate cancer. Further work will be needed to piece apart the impact of GSNO in the context in which it is applied, such as its spaciotemporal characteristics. The depth of the tumor microenvironment is vast, thus understanding how GSNO targets this three-dimensional space, albeit not individual cancer cells, would be of high importance. It is also interesting to note that GSNO bypasses the requirement for AR signaling in tumor xenografts, potentially pointing to its role to regulate other gene networks when functional AR is not available. Notable cell cycle genes, from our transcriptomic analysis, were significantly reduced by GSNO treatment both in-vivo and in-vitro, pointing to a correlation between AR signaling and cell cycle progression. We suggest a plausible model that could potentially explain how AR could exert a control on cell cycle targets through, for instance, promoter activity. In the presence of AR, FOXM1 promoter is being masked by cis-regulatory elements and/or AR itself, thus inhibiting genes involved in cell cycle and proliferation, while inhibition of AR releases the breaks on these targets, allowing for uncontrolled division to occur. This is one potential pitfall of long-term ADT, which makes AR obsolete while diminishing its important transcription factor properties. Instead, combinatorial therapies could be suggested with anti-proliferative and checkpoint agents that could aid ADT, while ensuring continued stepping on the breaks (**Figure 7**). Further work will be needed to test promoter and/or cis-regulatory element activity between AR and FOXM1 in the regulation of cell cycle proliferation. This research has the potential to propose new treatment interventions for patients with CRPC as we race to find a cure for this deadly and invasive disease.

**Figure 7.**
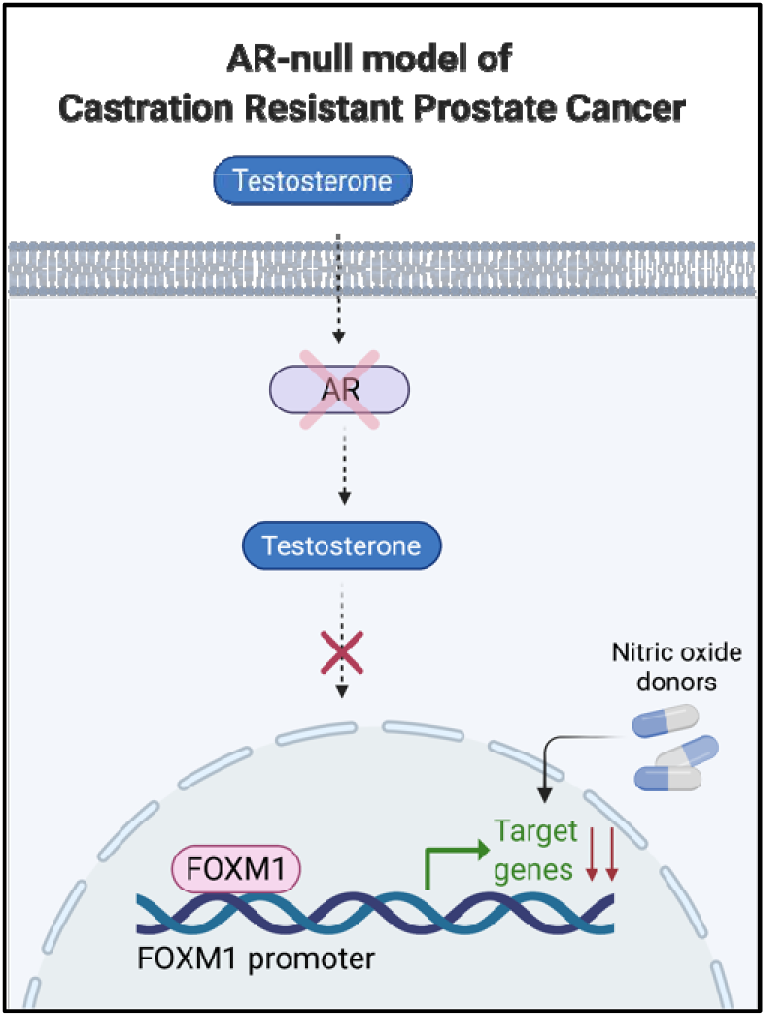
GSNO targets FOXM1 in castration resistant prostate cancer thereby inhibiting cell proliferation associated genes.

---

**Supplementary Figure 1.**
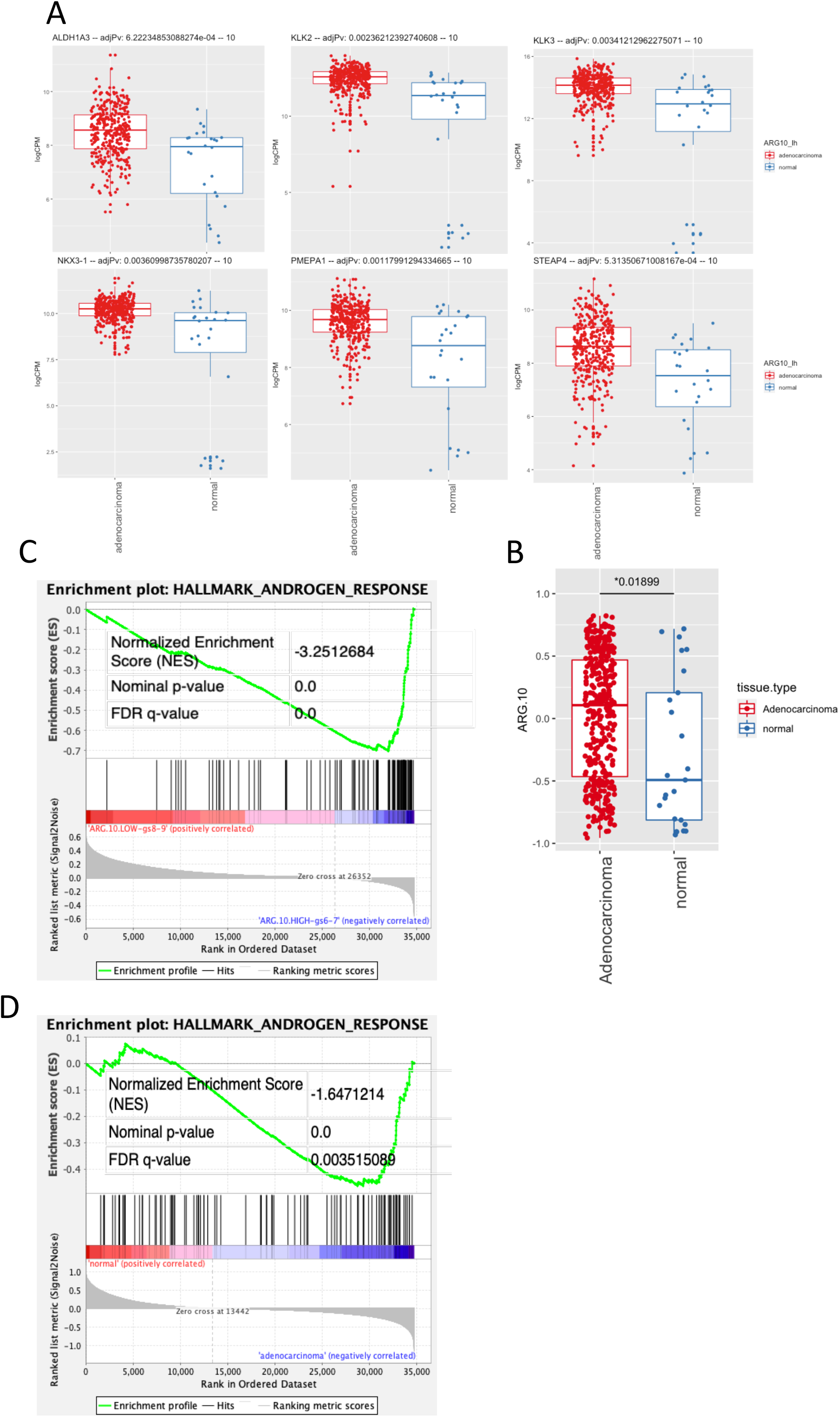
CRPC is associated with a reduction in AR activity in high-grade tumors. **A,** mRNA expression, normalized by logCPM, from RNA-seq data of Adenocarcinoma and normal patients in TCGA. **B,** gene set variation analysis score for ARG.10 in adenocarcinoma and normal patients (p=0.018, two tailed student’s t-test). **C,** gene set enrichment analysis plots for Hallmark Androgen Response in gleason8-9 and low ARG.10 score or gleason6-7 and high expression of ARG.10. **D,** additional enrichment for normal and Adenocarcinoma patients. NES, normalized enrichment score, FDR, false discovery rate.

**Supplementary Figure 2.**
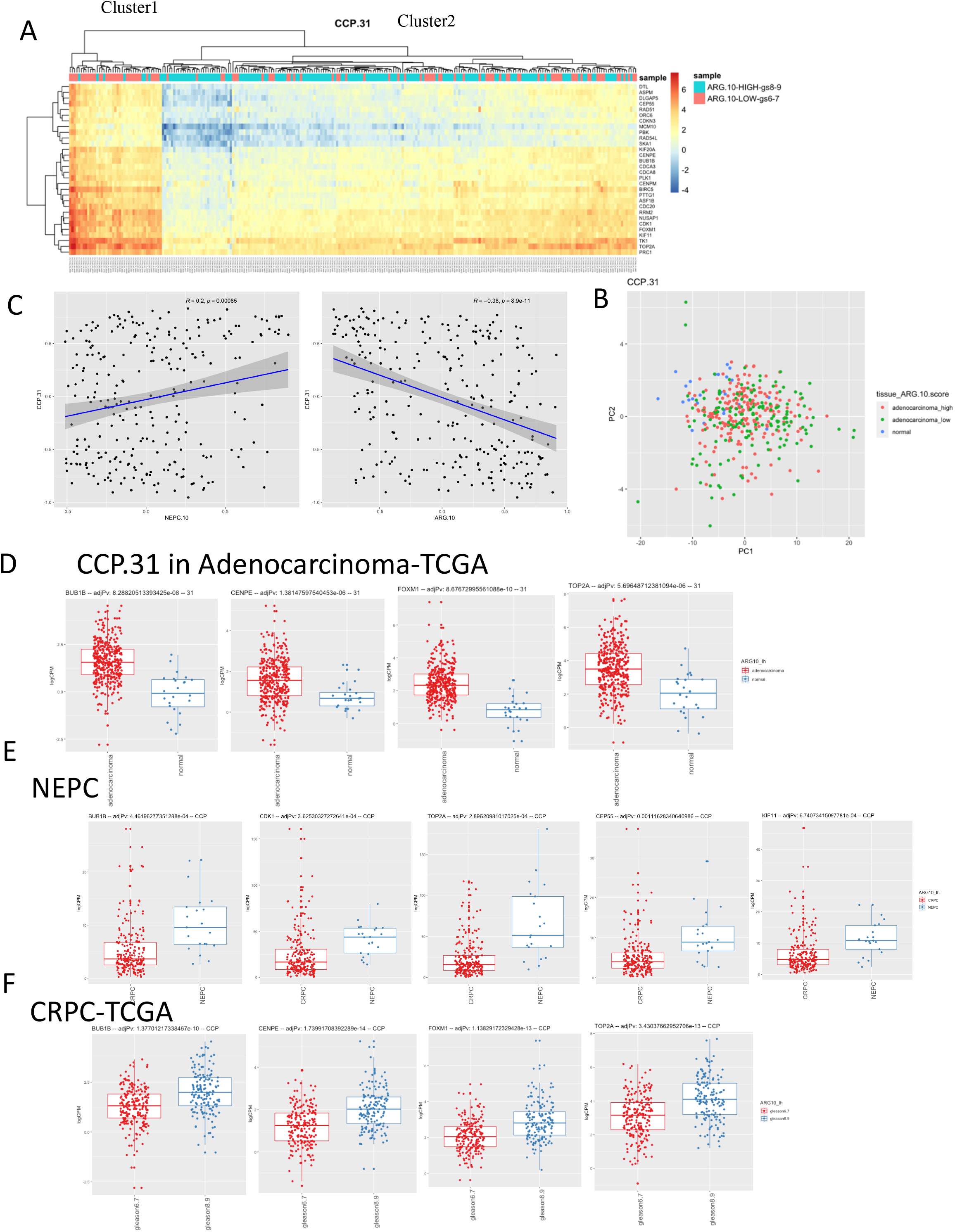
Transcriptomic analysis of high-grade, low AR-activity prostate tumors is enriched for cell cycle progression genes. **A,** Heatmap of unsupervised hierarchical clustering for CCP.31 genes depicting two distinct clusters enriched for corresponding sample groups annotated on the right of the diagram. **B,** principal components analysis plot, depicting distinct clustering of normal (blue) apart from Adenocarcinoma samples (red, green). **C,** Scatterplot of correlation between CCP.31 and ARG.10 (right) and CCP.31 and NEPC.10 (left) GSVA scores. Statistical significance is determined by regression. **D-F,** Tumor samples were grouped into high (AR.10>0; n=214) or low (AR.10<0; n=176) in TCGA and high (AR.10>0, n=148) or low (AR.10<0, n=120) in NEPC. **D,** boxplots of expression of representative genes reaching significance in CCP.31 list for indicated patient data sets. Expression is normalized and reported as log2-transformed counts per million. Statical significance (above the plots) is determined by ANOVA. P-values were adjustment by Bonferroni correction, calculated as alpha divided by the number of tests performed (first tier, number of targets within CCP pathway, second tier, number of patients in the cohort; n=268, NEPC; n=390, TCGA). Median expression indicated by thick middle line, first and third quartile by upper and lower line for TCGA, **D,** or NEPC, **E**. Adenocarcinoma patients were further divided into gleason8-9 to only include high-grade prostate cancer, **F**.

**Supplementary Table S1.**
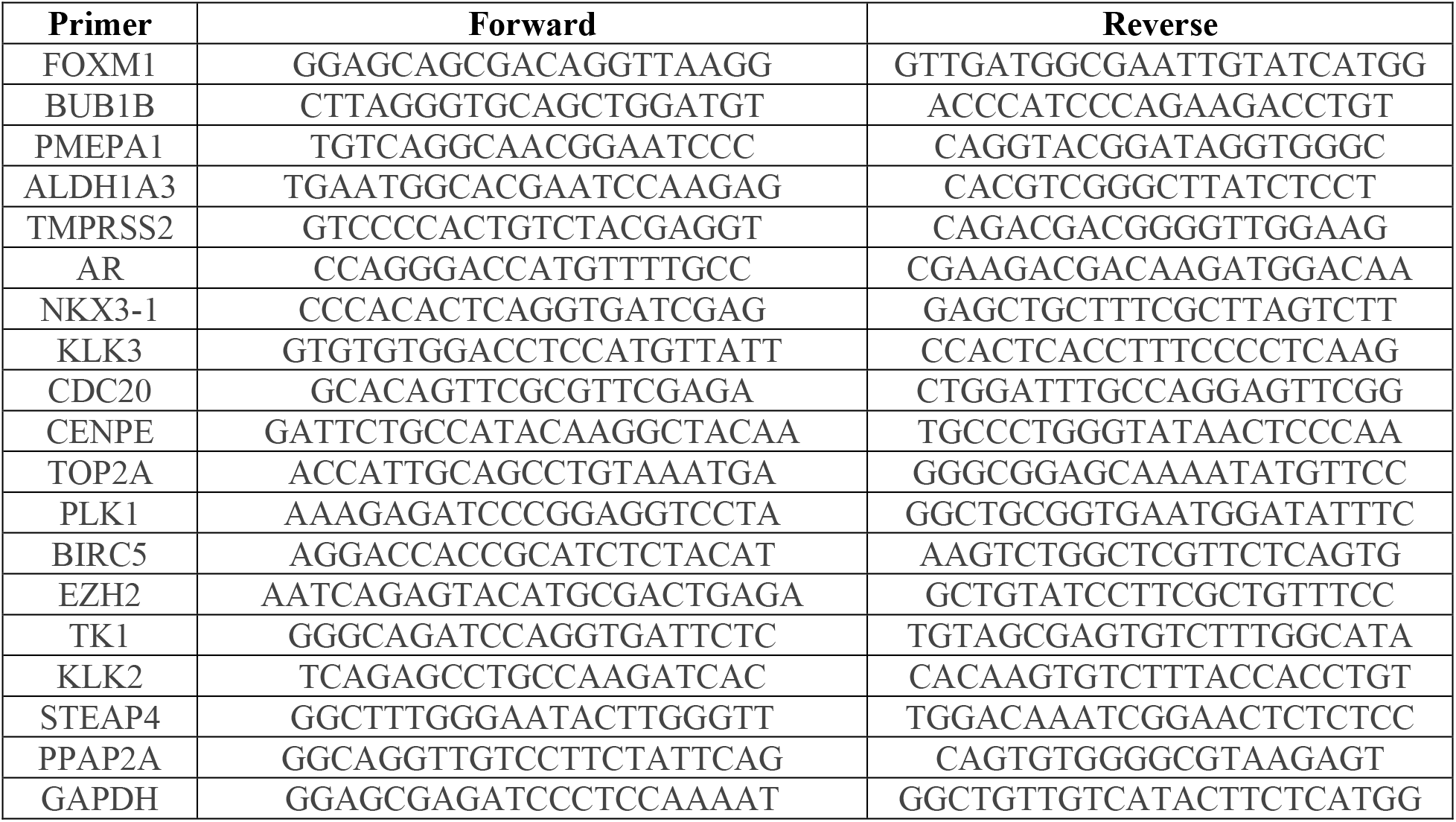
Complete list of primers used in RT-qPCR analysis on cell lysates and animal tumors.

## Notes

**Conflict of interest:** The authors declare no competing interests.

**Funding:** This work was supported by American Urological Association Research Scholar Award for HA, CTSI award for HA and RR, American Cancer Society, Stanley Glaser Award for RR, NIH grants to JMH 1R01 HL137355, 1R01 HL107110, 1R01 HL134558, 5R01 CA136387, and 5UM1 HL113460, and the Soffer Family Foundation.

### Competing Interest Statement

The authors have declared no competing interest.

## References

1. Jamroze A, Chatta G, Tang DG. Androgen receptor (AR) heterogeneity in prostate cancer and therapy resistance. Cancer Lett. 2021 Oct 10;518:1–9. doi: 10.1016/j.canlet.2021.06.006. Epub 2021 Jun 10. PMID: 34118355; PMCID: PMC8355210.

2. Li Q, Deng Q, Chao HP, Liu X, Lu Y, Lin K, Liu B, Tang GW, Zhang D, Tracz A, Jeter C, Rycaj K, Calhoun-Davis T, Huang J, Rubin MA, Beltran H, Shen J, Chatta G, Puzanov I, Mohler JL, Wang J, Zhao R, Kirk J, Chen X, Tang DG. Linking prostate cancer cell AR heterogeneity to distinct castration and enzalutamide responses. Nat Commun. 2018 Sep 6;9(1):3600. doi: 10.1038/s41467-018-06067-7. PMID: 30190514; PMCID: PMC6127155.

3. Bluemn EG, Coleman IM, Lucas JM, Coleman RT, Hernandez-Lopez S, Tharakan R, Bianchi-Frias D, Dumpit RF, Kaipainen A, Corella AN, Yang YC, Nyquist MD, Mostaghel E, Hsieh AC, Zhang X, Corey E, Brown LG, Nguyen HM, Pienta K, Ittmann M, Schweizer M, True LD, Wise D, Rennie PS, Vessella RL, Morrissey C, Nelson PS. Androgen Receptor Pathway-Independent Prostate Cancer Is Sustained through FGF Signaling. Cancer Cell. 2017 Oct 9;32(4):474–489.e6. doi: 10.1016/j.ccell.2017.09.003. PMID: 29017058; PMCID: PMC5750052.

4. Ketola K, Munuganti RSN, Davies A, Nip KM, Bishop JL, Zoubeidi A. Targeting Prostate Cancer Subtype 1 by Forkhead Box M1 Pathway Inhibition. Clin Cancer Res. 2017 Nov 15;23(22):6923–6933. doi: 10.1158/1078-0432.CCR-17-0901. Epub 2017 Sep 12. PMID: 28899970.

5. Nyquist MD, Ang LS, Corella A, Coleman IM, Meers MP, Christiani AJ, Pierce C, Janssens DH, Meade HE, Bose A, Brady L, Howard T, De Sarkar N, Frank SB, Dumpit RF, Dalton JT, Corey E, Plymate SR, Haffner MC, Mostaghel EA, Nelson PS. Selective androgen receptor modulators activate the canonical prostate cancer androgen receptor program and repress cancer growth. J Clin Invest. 2021 Jun 15;131(12):e151719. doi: 10.1172/JCI151719. Erratum for: J Clin Invest. 2021 May 17;131(10): PMID: 34128479; PMCID: PMC8203444.

6. Hickok JR, Thomas DD. Nitric oxide and cancer therapy: the emperor has NO clothes. Curr Pharm Des. 2010;16(4):381–91. doi: 10.2174/138161210790232149. PMID: 20236067; PMCID: PMC3782103.

7. Arora H, Panara K, Kuchakulla M, Kulandavelu S, Burnstein KL, Schally AV, Hare JM, Ramasamy R. Alterations of tumor microenvironment by nitric oxide impedes castrationresistant prostate cancer growth. Proc Natl Acad Sci U S A. 2018 Oct 30;115(44):11298–11303. doi: 10.1073/pnas.1812704115. Epub 2018 Oct 15. PMID: 30322928; PMCID: PMC6217394.

8. Liu Y, Gong Z, Sun L, Li X. FOXM1 and androgen receptor co-regulate CDC6 gene transcription and DNA replication in prostate cancer cells. Biochim Biophys Acta. 2014;1839(4):297–305. doi: 10.1016/j.bbagrm.2014.02.016. Epub 2014 Feb 27. PMID: 24583551.

9. Liu Y, Liu Y, Yuan B, Yin L, Peng Y, Yu X, Zhou W, Gong Z, Liu J, He L, Li X. FOXM1 promotes the progression of prostate cancer by regulating PSA gene transcription. Oncotarget. 2017 Mar 7;8(10):17027–17037. doi: 10.18632/oncotarget.15224. PMID: 28199985; PMCID: PMC5370019.

10. Kim MY, Jung AR, Kim GE, Yang J, Ha US, Hong SH, Choi YJ, Moon MH, Kim SW, Lee JY, Park YH. High FOXM1 expression is a prognostic marker for poor clinical outcomes in prostate cancer. J Cancer. 2019 Jan 1;10(3):749–756. doi: 10.7150/jca.28099. PMID: 30719174; PMCID: PMC6360432.

11. Beltran H, Prandi D, Mosquera JM, Benelli M, Puca L, Cyrta J, Marotz C, Giannopoulou E, Chakravarthi BV, Varambally S, Tomlins SA, Nanus DM, Tagawa ST, Van Allen EM, Elemento O, Sboner A, Garraway LA, Rubin MA, Demichelis F. Divergent clonal evolution of castration-resistant neuroendocrine prostate cancer. Nat Med. 2016 Mar;22(3):298–305. doi: 10.1038/nm.4045. Epub 2016 Feb 8. PMID: 26855148; PMCID: PMC4777652.

12. Robinson D, Van Allen EM, Wu YM, Schultz N, Lonigro RJ, Mosquera JM, Montgomery B, Taplin ME, Pritchard CC, Attard G, Beltran H, Abida W, Bradley RK, Vinson J, Cao X, Vats P, Kunju LP, Hussain M, Feng FY, Tomlins SA, Cooney KA, Smith DC, Brennan C, Siddiqui J, Mehra R, Chen Y, Rathkopf DE, Morris MJ, Solomon SB, Durack JC, Reuter VE, Gopalan A, Gao J, Loda M, Lis RT, Bowden M, Balk SP, Gaviola G, Sougnez C, Gupta M, Yu EY, Mostaghel EA, Cheng HH, Mulcahy H, True LD, Plymate SR, Dvinge H, Ferraldeschi R, Flohr P, Miranda S, Zafeiriou Z, Tunariu N, Mateo J, Perez-Lopez R, Demichelis F, Robinson BD, Sboner A, Schiffman M, Nanus DM, Tagawa ST, Sigaras A, Eng KW, Elemento O, Sboner A, Heath EI, Scher HI, Pienta KJ, Kantoff P, de Bono JS, Rubin MA, Nelson PS, Garraway LA, Sawyers CL, Chinnaiyan AM. Integrative Clinical Genomics of Advanced Prostate Cancer. Cell. 2015 Jul 16;162(2):454. doi: 10.1016/j.cell.2015.06.053. Epub 2015 Jul 16. PMID: 28843286.

